# Cigarette Smoke and Nicotine-Containing E-cigarette Vapor Downregulate Lung WWOX Expression Which is Associated with Increased Severity of Murine ARDS

**DOI:** 10.1101/2020.07.13.200832

**Authors:** Zhenguo Zeng, Weiguo Chen, Alexander Moshensky, Raheel Khan, Laura Crotty-Alexander, Lorraine B. Ware, C. Marcelo Aldaz, Jeffrey R. Jacobson, Steven M. Dudek, Viswanathan Natarajan, Roberto F. Machado, Sunit Singla

**Author notes:** Contributed equally to the work (Roberto F. Machado, Sunit Singla). Author contributions – Conception and design: SS, RFM, VN, SMD, JRJ, CMA, LBW, LCA; Data acquisition: ZZ, WC, AM, RK; Analysis and interpretation: SS, ZZ, WC, RFM; Manuscript preparation: SS, ZZ, RFM, CMA, LCA, SMD.

## Abstract

**RATIONALE:** A history of chronic cigarette smoking is known to increase risk for ARDS, but the corresponding risks associated with chronic e-cigarette use are largely unknown. The chromosomal fragile site gene, WWOX, is highly susceptible to genotoxic stress from environmental exposures, and thus an interesting candidate gene for the study of exposure-related lung disease.

**METHODS AND RESULTS:** Lungs harvested from current versus former/never smokers exhibited a 47% decrease in WWOX mRNA levels. Exposure to nicotine-containing e-cigarette vapor resulted in an average 57% decrease in WWOX mRNA levels relative to vehicle treated controls. In separate studies, endothelial (EC)-specific WWOX KO versus wild type mice were examined under ARDS-producing conditions. EC WWOX KO mice exhibited significantly greater levels of vascular leak and histologic lung injury. ECs were isolated from digested lungs of untreated EC WWOX KO mice using sorting by flow cytometry for CD31+CD45- cells. These were grown in culture, confirmed to be WWOX-deficient by RT-PCR and Western blotting, and analyzed by electric cell impedance sensing (ECIS) as well as a FITC dextran transwell assay for their barrier properties during MRSA or LPS exposure. WWOX KO ECs demonstrated significantly greater declines in barrier function relative to wild type cells during either MRSA or LPS treatment as measured by both ECIS and the transwell assay.

**CONCLUSION:** The increased risk for ARDS observed in chronic smokers may be mechanistically linked, at least in part, to lung WWOX downregulation, and this phenomenon may also manifest in the near future in chronic users of e-cigarettes.

## Introduction

Tobacco smoke exposure is the leading cause of preventable death worldwide due to its contribution to the pathogenesis of chronic cardiovascular and pulmonary diseases[1]. However, it has recently come to light that cigarette smoking likely also contributes to the development of an acute critical illness, the Acute Respiratory Distress Syndrome (ARDS)[2-6]. ARDS afflicts an estimated 200,000 patients/year in the United States alone, kills approximately 75,000, and is seriously debilitating for many survivors[7]. It occurs in select individuals following an inciting illness or event such as acute aspiration, pneumonia, sepsis, and blunt force trauma[8]. The cardinal, morbidity-producing feature of ARDS is non-cardiogenic pulmonary edema resulting from pulmonary vascular barrier disruption with consequent alveolar flooding, and respiratory failure[9, 10]. An increased incidence of ARDS has been observed in cigarette smokers versus nonsmokers following blunt force trauma, cardiac surgery, transfusion of blood products, or non-pulmonary sepsis[2-4, 6]. The mechanisms linking cigarette smoke (CS) exposure and ARDS development are largely unknown. It is hoped that research in this area may yield novel therapeutic targets for ARDS altogether, if not for a subset of patients in whom environmental exposures produce a mechanistic predisposition.

Recently, the link between CS exposure and ARDS risk has been highlighted further by the emergence of e-cigarette or vaping product use associated lung injury (EVALI) which has been occurring in epidemic proportions in the United States since the summer of 2019[11]. A significant proportion of EVALI seen thus far appears to be associated with specific chemical additives that have been vaped[12, 13]. The overall rise in use of e-cigarettes has been fueled, in part, by the unsubstantiated notion that they are safer than traditional cigarettes because of the absence of combustion products in e-cigarette vapor versus CS. Thus, they are being used in many cases as a strategy for patients to quit smoking traditional cigarettes[14]. However, notwithstanding the specific circumstances underlying the current EVALI epidemic, very little, if anything, is known about the long-term consequences of e-cigarette vapor exposure in terms of ARDS risk, as well as risks for the development of other CS-associated lung diseases.

A multitude of epigenetic modifications caused by cigarette smoke, any combination of which may precipitate the CS-ARDS connection, include methylation at various CpG-containing sequences[15], to more dramatic changes such as rearrangements and deletions that occur with some frequency at chromosomal fragile sites[16, 17]. Amongst both of these categories of CS-induced gene changes is the tumor suppressor, WWOX, which resides at the second most active chromosomal fragile site in the human genome[18, 19]. Apart from its heightened susceptibility to loss of function mutations and methylation induced downregulation via environmental exposures including cigarette smoke, its potential role in the pathogenesis of lung disease was elucidated by gene knockdown studies in mice and cultured pulmonary epithelial cells[20]. Acute si-RNA mediated knockdown of lung WWOX expression in mice was observed to result in neutrophilic inflammation, a finding that may be relevant to the pathogenesis of chronic lung diseases such as COPD[20]. An interesting finding in this study was the presence of increased LPS-induced pulmonary vascular leak during lung WWOX knockdown that was out of proportion to the degree of neutrophilic inflammation[20]. This raised the question of whether an inflammation-independent mechanism of endothelial barrier vulnerability was a part of the WWOX-deficient phenotype.

In the current study, specific endothelial silencing of WWOX expression in cultured cells and mice was achieved to assess WWOX as a candidate gene for CS-ARDS.

## Materials and Methods

### Reagents

FITC-Dextran, SP100625, PP2, LPS, RBC Lysis Buffer, DMEM, bovine serum albumin, DNase, Dispase II, and RIPA Buffer were purchased from Sigma-Aldrich. BCA Protein Assay Kit, Shandon Kwik-Diff kit, siPORT Amine transfection reagent, Lipofectamine 2000, High Capacity cDNA Reverse Transcription kit, Taqman Gene Expression Master Mix, and Taqman Gene Expression Assays for GAPDH and WWOX were from Thermo Fisher. Mini-PROTEAN® TGX™ Precast Gels were from BioRad. QIAShredder and RNeasy Mini kits were from Qiagen. Anti-WWOX antibody was from abcam (Cambridge, MA). Human lung microvascular endothelial cells (ECs), and endothelial cell growth media were purchased from Lonza (Basel, Switzerland). Heparin was purchased from Hospira (Lake Forest, IL). Collagenase was from Worthington (Lakewood, NJ). ELISA kits for mouse cytokines (KC, MIP-2, MCP-1, IL-6, IL-1β, TNF-α), anti-mouse CD16/32, anti-mouse CD31-PE/Cy7, and anti-mouse CD45-Alexa700 antibodies were from BioLegend (San Diego, CA) and R&D systems (Minneapolis, MN). siGENOME WWOX-targeting and scrambled control siRNA was from Dharmacon/GE. pCMV entry vector containing the ORF corresponding to (Myc-DDK-tagged)-human WW domain-containing oxidoreductase ((WWOX), transcript variant 1, as well as a (Myc-DDK-tagged)-empty vector were purchased from OriGene.

### Cell culture

ECs obtained from Lonza were grown in cell culture using endothelial growth media containing supplemental growth factors and 2% fetal bovine serum according to the manufacturer’s instructions. Cells underwent a maximum of three passages for use in experiments.

For separate studies, ECs were isolated from the lungs of 8 week old C57BL/6-WWOXflox/flox and Cdh5-CreERT2/WWOX^flox/flox^ mice as in Kawasaki et al[21]. Following ketamine/xylazine administration, the thoracic cavity was surgically opened. 20 ml of sterile PBS containing 10U/ml heparin was instilled into the pulmonary vasculature via injection through the right ventricle until the lungs were blood-free. Mouse lungs were then harvested, minced, and digested in an enzyme complex as previously described. Washed cell pellets were resuspended and serially incubated with anti-mouse CD16/32 (to block Fc receptors), anti-mouse CD31-PE/Cy7, and anti-mouse CD45-Alexa700 antibodies. After washing with PBS/BSA, cells were sorted by flow cytometry (MoFlo Astrios Cell Sorter, Beckman Coulter, Indianapolis, IN) to isolate CD31+CD45-ECs which were then placed in EC growth media on gelatin coated plates.

### USA 300 MRSA

Wild-type USA 300 MRSA was prepared as in Chen et al[22]. Briefly, 3 ml of tryptic soy broth was inoculated with a single colony of bacteria and incubated overnight. 1 ml of this overnight culture was then used to inoculate 100 ml of tryptic soy broth for incubation until and optical density of 0.5 at 660 nm was reached. This was then centrifuged and washed for resuspension in sterile PBS at a concentration of 1 × 10^8^ CFU per 30 µl. MRSA was heat-killed by incubation at 65°C for 1 hour prior to use in cell-based experiments. Live MRSA was used in mice.

### Animals

All experiments and animal care procedures were approved by the University of Illinois at Chicago Animal Care and Use Committee. C57BL/6-WWOX^flox/flox^ mice were donated generously by Dr. C. Marcelo Aldaz, MD/PhD of MD Anderson Center at the University of Texas. C57BL/6-Cdh5-CreERT2 mice were purchased from Jackson Laboratories. The two strains were crossed to produce Cdh5-CreERT2/WWOX^flox/flox^ mice as in Ludes-Meyers et al[23]. They were housed in cages in a temperature controlled room with a 12 hr dark/light cycle, and with free access to food and water.

### E-cigarette exposure in Mice

6-8 week old male C57Bl/6 mice were exposed to vehicle versus nicotine-containing e-cigarette vapor via inExpose inhalation system (Scireq) for 1 hour a day, 5 days a week, and for 3 months. Thirty to sixty minutes after the final exposure, mice were anesthetized with ketamine (100mg/kg) and xylazine (10mg/kg), and lungs were harvested as described below. All e-cigarette related animal protocols were approved by the University of California, San Diego IACUC Committee.

### Murine models of ARDS

6 week old male Cdh5-CreERT2/WWOX^flox/flox^ mice received intraperitoneal injections of tamoxifen (75mg/kg) for 5 consecutive days. Control animals received an equivalent volume of vehicle (corn oil). 7 days later they were anesthetized with isoflurane and underwent nasal instillation of USA 300 MRSA (0.75 × 10^8^ CFU) versus an equivalent volume of sterile PBS. Mice were then housed in cages as described above for 18 hours after which they underwent testing and lung harvesting as described below. In separate experiments, tamoxifen versus vehicle treated mice were anesthetized with isoflurane and underwent intratracheal instillation of 1mg/kg LPS versus an equivalent volume of sterile saline. 18 hours later they underwent testing and lung harvesting as described below.

### FITC-Dextran permeability assay

70 kDa FITC-Dextran was used to measure pulmonary vascular leak as described previously [24]. Briefly, FITC-Dextran was mixed to a concentration of 30 mg/ml in sterile PBS and sterile filtered. 30 minutes prior to euthanasia, 100 µl of FITC-Dextran solution was injected into the retro-orbital venous plexus. Bronchoalveolar lavage fluid (BALF) was collected as described below. Blood was collected during lung harvest and centrifuged to isolate serum. Fluorescence of equal volumes of BALF supernatant and serum were measured and relative changes in permeability were expressed as a ratio of FluoBALF/FluoSerum normalized to the control condition.

### Bronchoalveolar lavage fluid (BALF) collection and analysis

Mice were anesthetized with ketamine/xylazine. The trachea was exposed surgically and a small incision was made on the anterior surface. An 18-gauge blunt-end cannula was inserted into this opening and secured with a suture tied around the trachea. 1 ml of sterile saline was infused through this cannula and slowly withdrawn.

This BALF was centrifuged to pellet cells and remove supernatant. RBCs were removed from the cell pellet using RBC lysis buffer according to the manufacturer’s instructions. The remaining cells (leukocytes) were pelleted, resuspended in a fixed volume of saline, and a small aliquot was used to measure cell concentration with an automated cell counter. The remaining cell suspension was used to make a cytospin prep on a glass slide for staining with the Shandon Kwik-Diff kit. A manual differential cell count was performed on 10 hpfs to determine relative percentages of neutrophils in the leukocyte population. Protein concentration was measured using the BCA Protein Assay kit, and the corresponding ELISA kits were used to measure concentrations of KC, MIP-2, MCP-1, IL-6, IL-1β, TNF-α.

### Lung harvesting for RT-PCR, Western blotting, and Histology

Following BALF collection, the right ventricle was infused with sterile saline until the lungs were free of intravascular blood. The right middle lobe was excised and snap frozen in liquid nitrogen for RT-PCR, and Western blotting. The remaining lobes were excised and placed in 10% formalin for 24 hours followed by transfer to 70% ethanol for storage until paraffin embedding and sectioning for H&E staining.

For RT-PCR, lungs were thawed, lysed, homogenized, and RNA was isolated using the QIAShredder and RNeasy Mini kits per manufacturer’s instructions. cDNA was produced for each sample using the high capacity cDNA Reverse Transcription kit from Thermo Fisher. Real-time PCR was performed using the Taqman Gene Expression Master Mix and Assays per manufacturer’s instructions. Relative quantification between controls and experimental samples of WWOX gene expression normalized for GAPDH was determined using the ΔΔC_t_ method.

For Western blotting, snap frozen lungs were thawed and homogenized in RIPA buffer. Homogenized samples were centrifuged to pellet debris and isolate protein lysates. Protein concentration of lysates was measured using the BCA Protein Assay kit, and samples were diluted in RIPA buffer until all had the same protein concentration. After dilution with SDS buffer, samples were loaded into 4-20% Mini-PROTEAN® TGX™ Precast Gels and subjected to SDS-PAGE. Gels were transferred to nitrocellulose membranes which were then subjected to Western blotting using antibodies targeting proteins of interest in accordance with the manufacturer’s instructions. Densitometric analysis was performed using ImageJ software.

### Electric Cell Impedance Sensing (ECIS) Permeability Assay

Endothelial cells were grown to confluence in polycarbonate wells containing evaporated gold microelectrodes purchased from Applied BioPhysics and measurements of transendothelial electrical resistance (TER) were performed using an ECIS system (Applied BioPhysics) as described previously[25-27]. Briefly, a 4000-Hz alternating current was applied across the electrodes with a fixed amplitude and external resistance to establish a constant current of approximately 1 µA. The in-phase and out-of-phase voltages between the electrodes were measured in real time and were used to calculate transendothelial resistance (TER). Increased cell adherence and confluence correlates with higher TER, while cell retraction, rounding, or loss of adhesion leads to decrease in TER[28]. TER was monitored for stability prior to initiating MRSA/LPS treatment in all experiments and a threshold of 2.5 x 10^3^Ω was required for ECs to be considered confluent on the electrodes and suitable for study. TER from each microelectrode was measured at discrete time intervals and recorded for analysis.

### FITC-dextran Transwell Permeability Assay

ECs were grown to confluency on transwell inserts (EMD Millipore) containing 3 micrometer pores. 10 μg/ml FITC-labeled dextran (70 kDa) was added to the media over the cells along with either vehicle, heat killed MRSA (2 × 10^8^CFU/ml), or LPS (1µg/ml). After 2 hours, 100 μl of media was extracted from below the transwell insert for measurement of FITC concentration by fluorometry (excitation/emission spectrum peak wavelengths of 495 nm and 519 nm) to determine the degree of EC monolayer permeability as described previously[29].

### In vitro silencing of WWOX in human pulmonary endothelial cells

In vitro silencing of WWOX in human ECs (Lonza) was achieved using transfection of siRNA as described in Wolfson, et al[30]. WWOX-targeting (CCAAGGACGGCUGGGUUUA) versus scrambled control siRNA (UAGCGACUAAACACAUCAA) was complexed with siPORT Amine transfection reagent. Serum free media was used to bring the concentration of siRNA to 100 nM. Transfection of ECs was then performed as described previously[30]. 72 hours after transfection, ECs were analyzed by Western blotting for WWOX expression, and by ECIS for barrier function during LPS exposure.

### Human Lung Tissue

Vanderbilt IRB approved this work as non-human subjects research (ie no IRB approval required). Lung tissue specimens were procured from organ donors whose lungs were declined for transplantation as part of the Beta-agonist for Oxygenation in Lung Donors study[31]. At the time of procurement, lungs were resected without perfusion and were transported on ice to the investigators’ laboratory. Portions of each lobe were sampled and immediately frozen at -80C in RNALater (Qiagen) until RNA extraction. Clinical history including history of lung disease and cigarette smoking was obtained from the donor medical record.

### Statistical analysis

Either a Student’s t-test or an ANOVA were performed where applicable on all data using GraphPad Prism software. *p<0.05.

## Results

### Lung tissue from current human smokers versus former/never smokers exhibits decreased WWOX mRNA expression

RNA was extracted from human lung tissue samples taken from lung donors with a history of regular cigarette smoking and analyzed by RT-PCR for levels of WWOX mRNA expression compared to that of former or never smokers. As shown in **Figure 1A**, lung WWOX expression was 47% lower in current versus former or never smokers (p<0.05).

**Figure 1.**
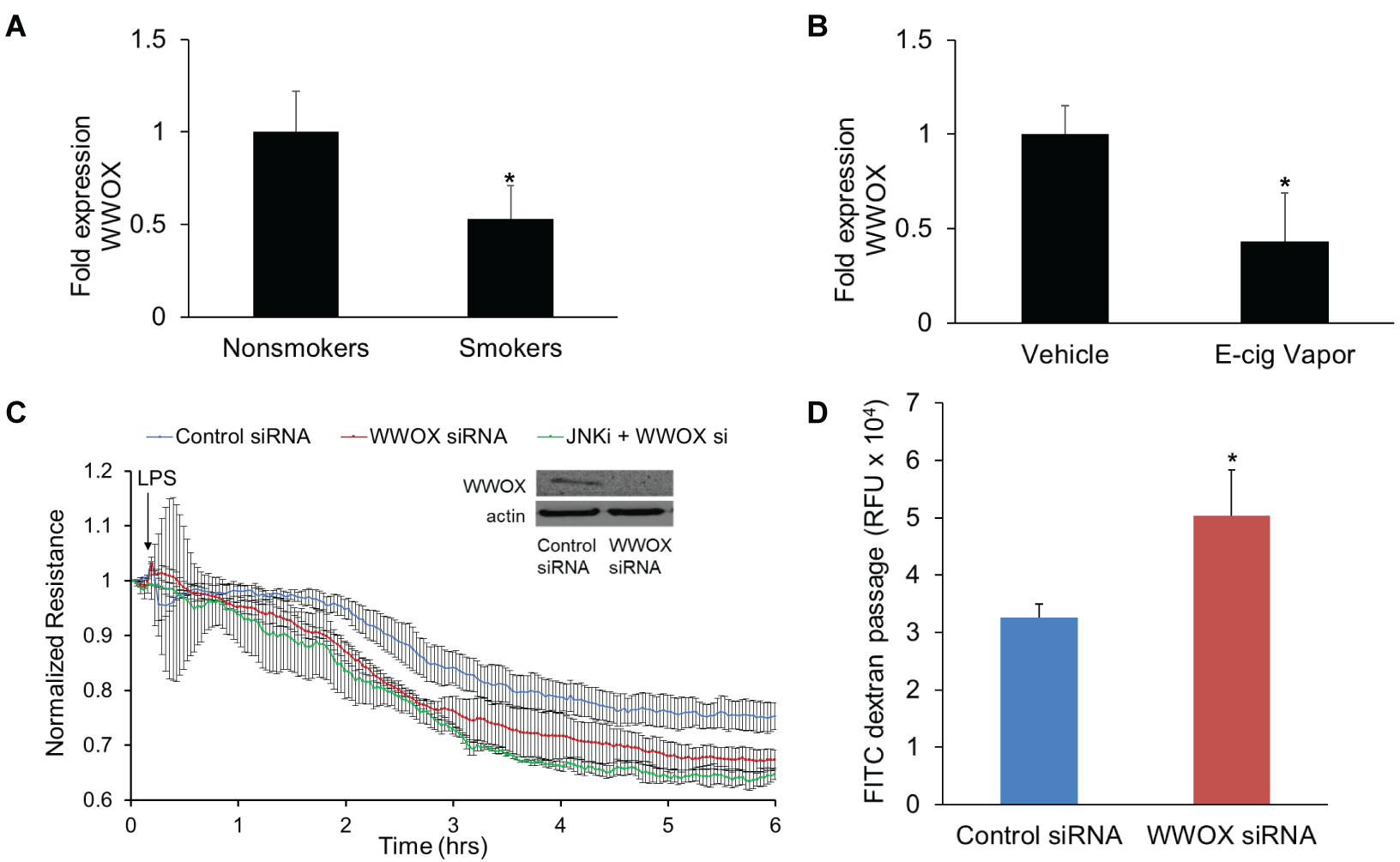
WWOX expression in human smokers/e-cigarette-exposed mice, and the associated lung endothelial barrier consequences. **A.** RNA was extracted from lung tissue harvested from current versus never/former human smokers. RT-PCR for WWOX (Taqman® Assay, Life Technologies; Carlsbad, CA) was performed. The bar graph depicts fold-changes in WWOX mRNA expression calculated by the ΔΔC_t_ method with GAPDH as the internal control. Results are shown as means +/- SD in n=3 independent experiments. **B.** RNA was extracted from lung tissue harvested from mice exposed to e-cigarette vapor with or without nicotine (control). RT-PCR for WWOX results are shown here. Results are shown as means +/- SD in n=3 independent experiments. **C.** Pulmonary endothelial cells (ECs) were transfected with either a WWOX-targeted or control scrambled siRNA (Dharmacon; Lafayette, CO). A western blot for WWOX confirms silencing. Transfected ECs were grown to confluency on gold electrodes for electrical cell impedance sensing (ECIS) analysis. They were pretreated with a pharmacologic inhibitor of JNK (SP600125) versus vehicle followed by LPS. **D.** In separate experiments, ECs were grown in a monolayer on transwell inserts containing 3 micron pores. 10 µg/ml FITC-labeled dextran (3kDa) was added over the cells along with LPS. After 1 hour media from below the transwell was analyzed by fluorometry for the presence of FITC. Results are shown as means +/- SD in n=3 independent experiments. *p<0.05, compared with control by Student’s t-test.

### Nicotine-containing e-cigarette vapor exposure in mice causes knockdown of mouse lung WWOX expression

C57Bl/6 mice were exposed to vehicle versus nicotine-containing e-cigarette vapor for 3 months. Lungs were harvested and examined by RT-PCR for levels of WWOX expression. Lungs from mice exposed to nicotine-containing e-cig vapor exhibited an average 57% decrease in WWOX expression compared to controls (p<0.05) (**Figure 1B**).

### WWOX-silenced human pulmonary endothelial cells sustain greater barrier dysfunction during LPS exposure

Human lung microvascular endothelial cells (hLMVECs) were treated with scrambled, control versus WWOX-targeting siRNA which resulted in efficient knockdown of WWOX expression (**inset, Figure 1C**). Cells were then grown to confluency on gold electrodes for ECIS assessments of TER in response to LPS. As shown in **Figure 1C**, treatment of WWOX-silenced hLMVECs with LPS resulted in larger decreases in TER compared to wild-type cells. In an earlier study[20], WWOX knockdown was associated with JNK-dependent pro-inflammatory signaling in alveolar epithelial cells. Therefore, WWOX-silenced hLMVECs were pretreated with the JNK inhibitor, SP600125, prior to LPS exposure to determine whether a JNK-related pathway was involved in heightened barrier dysfunction related to WWOX knockdown in endothelial cells. JNK inhibition prior to LPS resulted in no significant rescue of barrier function towards that of wild type cells (**Figure 1C**).

To confirm that the observed decreases in TER in control versus WWOX-silenced hLMVECs during LPS exposure correlated with increased permeability of the EC monolayer, trans-monolayer FITC-dextran flux was measured. Control versus WWOX-silenced hLMVECs were grown to confluency on transwell inserts containing 3 µm pores, and then treated with 10 µg/ml FITC-dextran (3kD) added to the media overlying the monolayer. After exposure to 100 ng/ml LPS for 2 hours, media from the chamber underlying the transwell inserts was extracted for fluorometry. Knockdown of WWOX expression resulted in increased LPS-induced trans-monolayer flux of FITC-dextran relative to wild type monolayers (**Figure 1D**).

### Generation of Endothelial-specific WWOX KO mice

C57BL/6-WWOX^flox/flox^ mice were bred with C57BL/6-Cdh5-CreERT2 to generate tamoxifen-inducible EC-specific WWOX KO mice. Cre+ flox homozygotes were confirmed by PCR as in Ludes-Meyers et al[23] (**Figure 2A)**. Following tamoxifen treatment in mature animals to induce endothelial-specific WWOX knockdown, lungs were harvested for flow cytometric isolation of CD31+CD45-endothelial cells (**Figure 2B**). Cells were propagated in culture and analyzed by RT-PCR (**Figure 2C**) and Western blotting (**Figure 2D**) to confirm WWOX knockdown.

**Figure 2.**
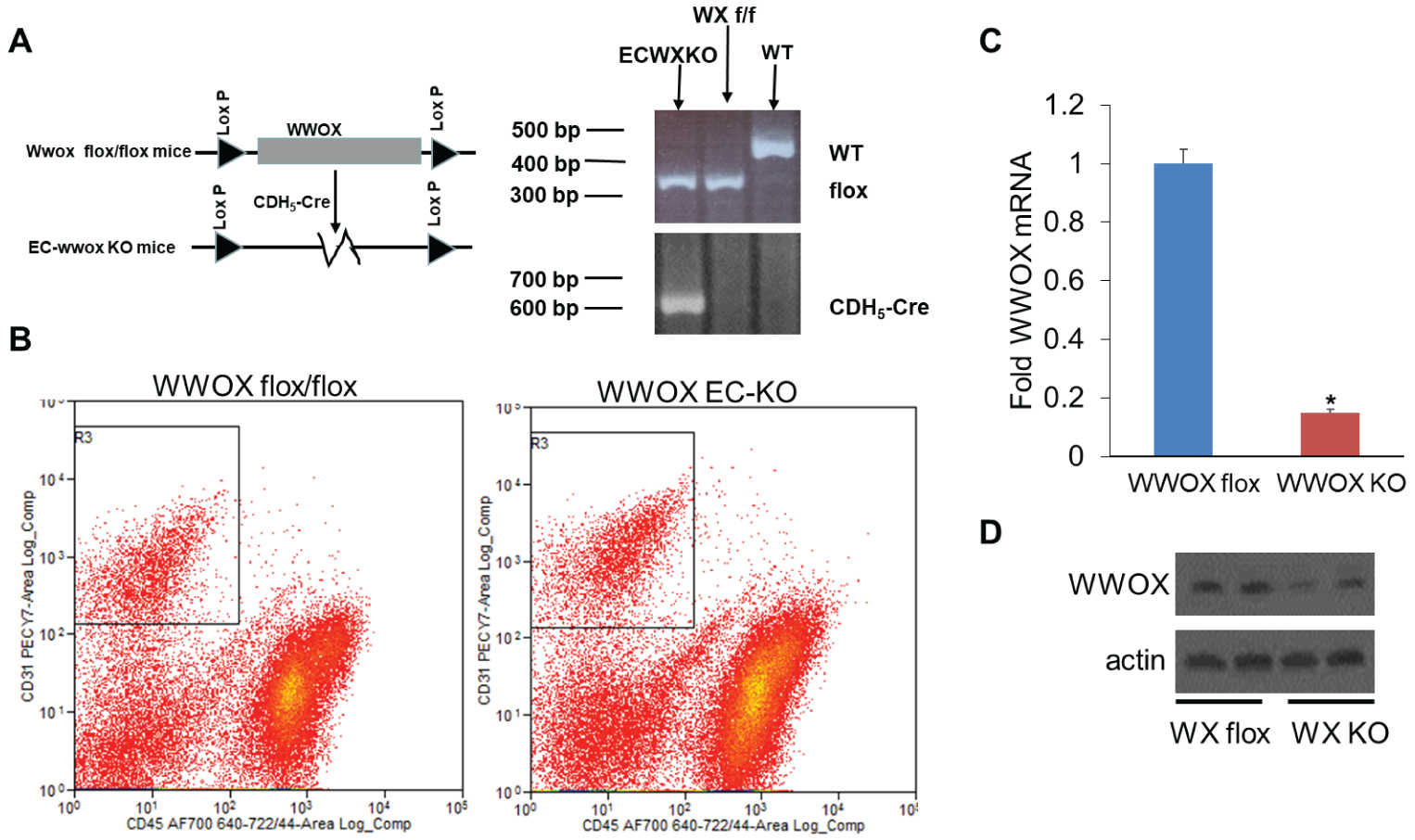
Generation of inducible endothelial-specific WWOX KO mice. WWOX flox/flox mice were obtained from the lab of Dr. C. Marcelo Aldaz, MD/PhD of MD Anderson Cancer Center at the University of Texas. These were bred with mice containing a tamoxifen-inducible cre deleter under the control of the CDH5 promoter to obtain an inducible endothelial-specific WWOX KO. **A.** Genotyping was performed as described in Ludes-Meyers, et al. PLoS One. 2009; 4(11): e7775. **B.** Lung endothelial cells were isolated from cell suspensions obtained from WWOX flox and KO mice using FACS to segregate CD45-CD31+ cells. These were grown in culture and analyzed for WWOX expression by RT-PCR (**C**) and Western Blotting (**D**). Results are shown as means +/- SD in n=3 independent experiments. *p<0.05, compared with control by Student’s t-test.

### EC-specific WWOX KO mice exhibit greater amounts of pulmonary vascular leak following intratracheal LPS instillation

A direct injury model of ARDS via intratracheal instillation of LPS was used to phenotype differences in vascular leak between EC WWOX KO mice and wild type controls. As shown in **Figure 3**, bronchoalveolar lavage fluid (BALF) obtained from EC WWOX KO mice treated with LPS exhibited greater amounts of leukocyte numbers, protein concentration, and FITC-dextran flux from the vascular space compared to LPS-treated wild type mice.

**Figure 3.**
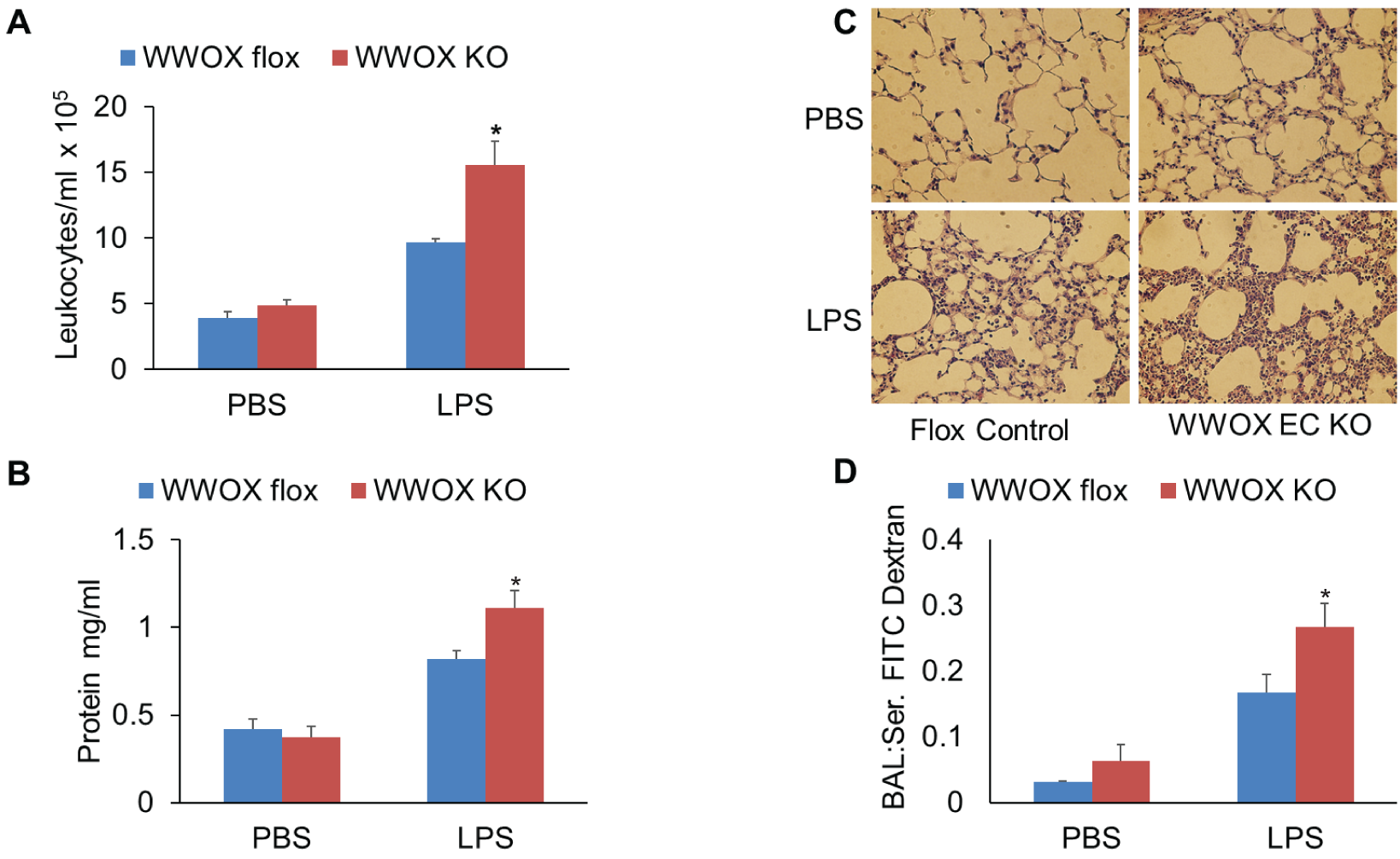
EC-specific WWOX KO mice exhibit greater vascular leak during LPS-induced ARDS. 6 WWOX flox (control) and 6 WWOX KO mice were treated with intratracheal instillation of either PBS or 1 mg/kg LPS (n=3 per group). Eighteen hours later all mice underwent bronchoalveolar lavage (BAL) using 1 ml of PBS. Lungs were then harvested for Western blotting and histologic examination. **A, B, D**: Bar graphs depict BAL leukocyte counts, protein concentration, and FITC dextran flux as a measure of permeability. Results are shown as means +/- SD in n=3 independent experiments. *p<0.05, compared with LPS-treated control by Student’s t-test. **C.** A representative hematoxylin-eosin-stained lung histologic section from each of the 4 experimental conditions is shown here.

Unlike the previously reported observations in whole lung WWOX knockdown mice[20], there were no significant differences between untreated EC WWOX KO mice and controls. Increased vascular leak of cells and protein in LPS-treated EC WWOX KO mice was confirmed by histologic examination of H & E stained lung sections (**Figure 3D**).

Interestingly, while there was a trend towards increased concentrations of inflammatory cytokines in the BALF of EC WWOX KO mice treated with LPS compared to wild types, none were statistically significant (**Figure 4**). This, along with the ECIS observations in **Figure 1**, which showed no rescue effect of JNK inhibition in the enhanced vascular leak noted in WWOX-silenced ECs, suggests an inflammation-independent EC-specific mechanism underlies these phenomena.

**Figure 4.**
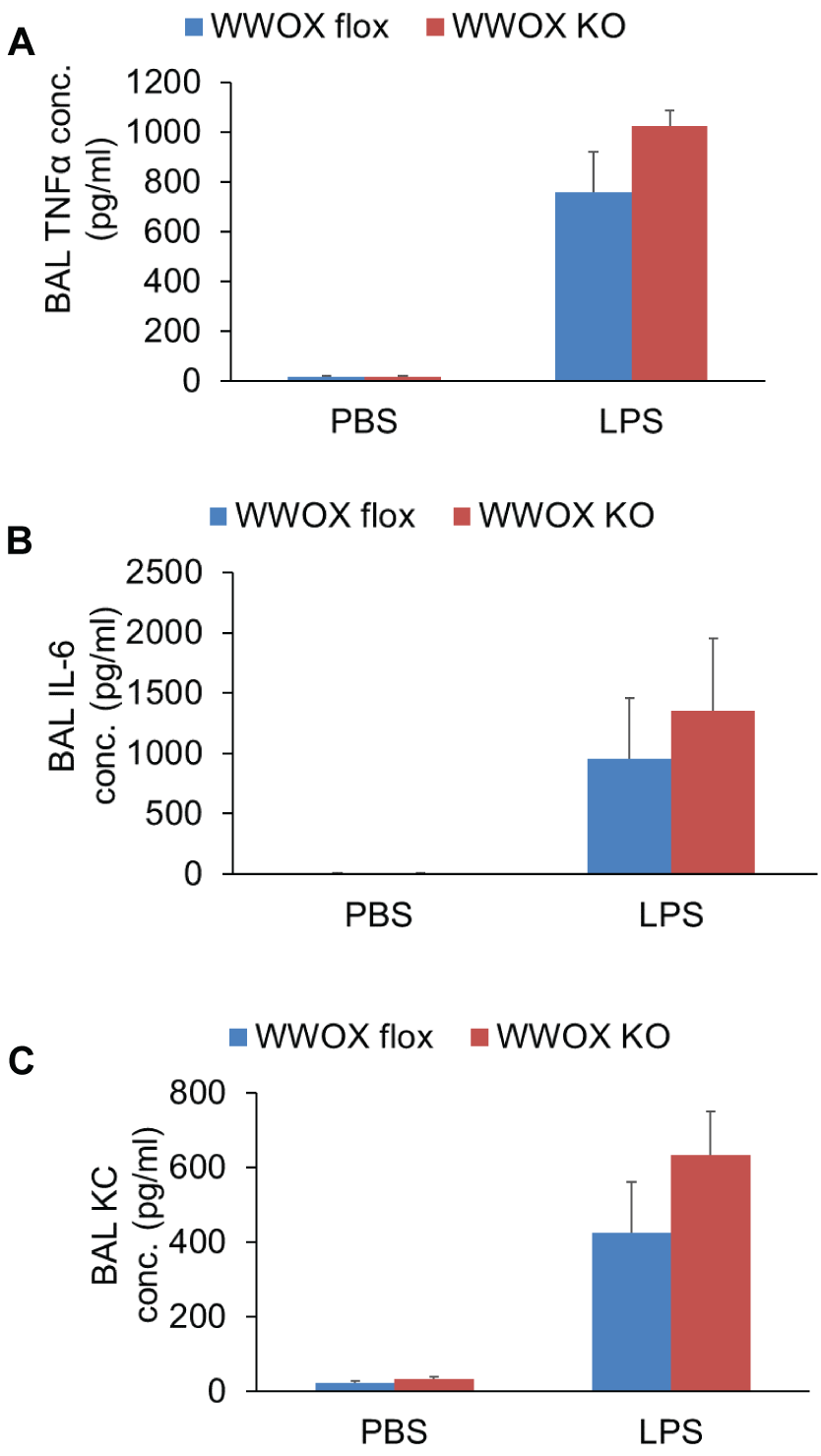
EC-specific WWOX KO mice do not exhibit greater BAL cytokines during LPS-induced ARDS. Bar graphs depict BAL TNFα, IL-6, and keratinocyte-derived chemokine (KC) concentrations. None of the increases in LPS-treated WWOX KO animals were statistically significantly different than those of LPS-treated controls. Results are shown as means +/- SD in n=3 independent experiments. *p<0.05, compared with LPS-treated control by Student’s t-test.

### EC-specific WWOX KO mice injured in a whole bacterial MRSA model of ARDS exhibit greater amounts of vascular leak without significant increases in inflammation compared to wild type mice

To further test the notion that enhanced vascular leak observed in EC WWOX KO mice is an inflammation-independent phenomenon, and to extend these observations to a more common clinical scenario, a whole bacterial model of ARDS induced by intratracheal instillation of heat-killed USA 300 MRSA was used to further phenotype these animals. As shown in **Figures 5A and 5B**, BALF from EC WWOX KO animals treated with MRSA exhibited greater amounts of leukocyte numbers and protein concentration compared to corresponding MRSA-treated wild types. Histologic examination of H & E stained lung sections confirmed increased vascular leak in EC WWOX KO mice (**Figure 5C**).

**Figure 5.**
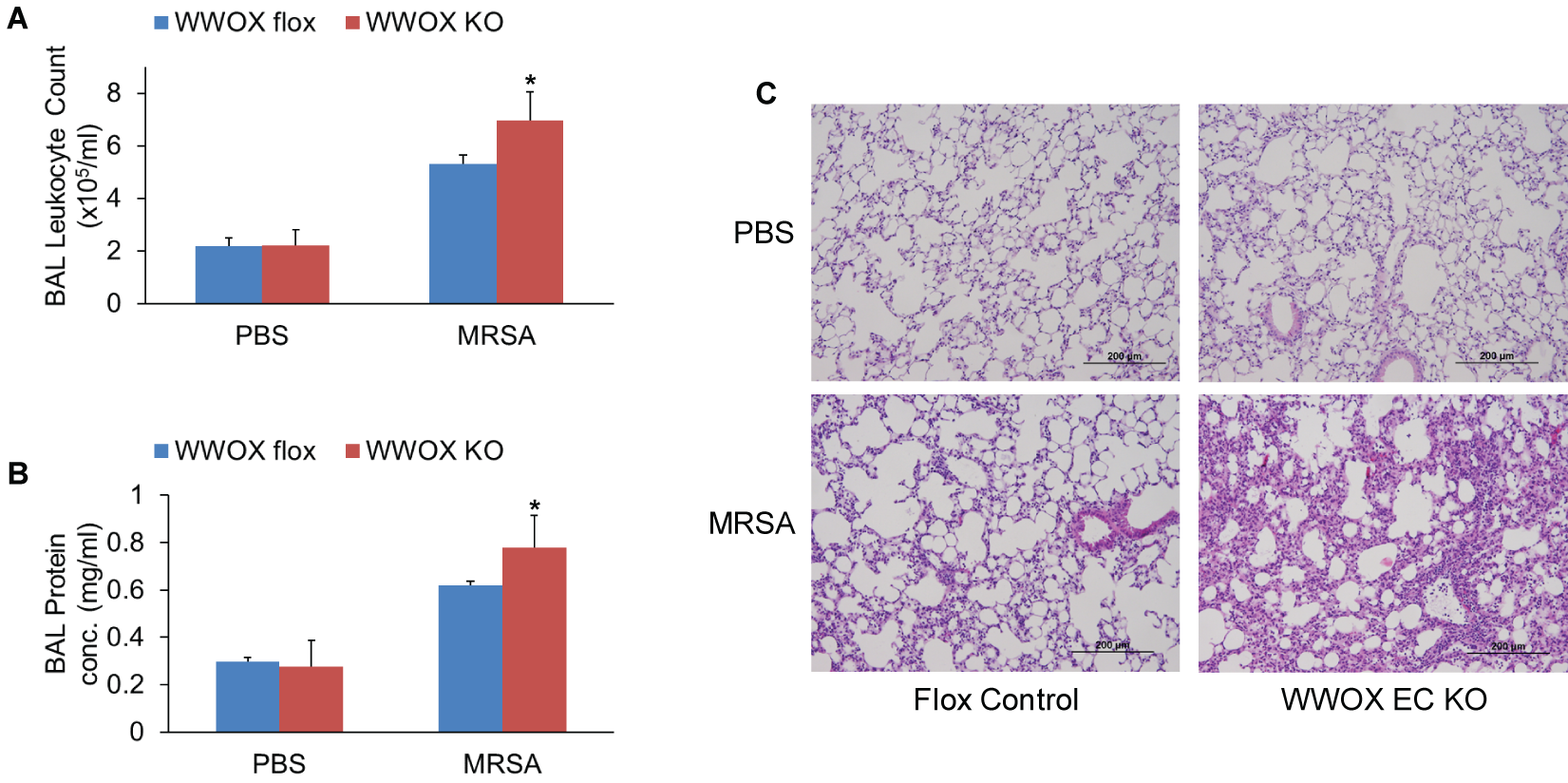
EC-specific WWOX KO mice exhibit greater vascular leak during MRSA-induced ARDS. 6 WWOX flox (control) and 6 WWOX KO mice were treated with intratracheal instillation of either PBS or 1 × 10^8^ CFU of USA 300 MRSA (n=3 per group). Eighteen hours later all mice underwent bronchoalveolar lavage (BAL) using 1 ml of PBS. Lungs were then harvested for Western blotting and histologic examination. **A, B**: Bar graphs depict BAL leukocyte counts, and protein concentration as a measure of permeability. Results are shown as means +/- SD in n=3 independent experiments. *p<0.05, compared with LPS-treated control by Student’s t-test. **C.** A representative hematoxylin-eosin-stained lung histologic section from each of the 4 experimental conditions is shown here.

Interestingly, once again while there was a trend towards increased proinflammatory cytokine (TNFα, IL-6, KC) secretion in BALF from MRSA-treated EC WWOX KO animals versus wild type counterparts, these differences were not statistically significant (**Figure 6**).

**Figure 6.**
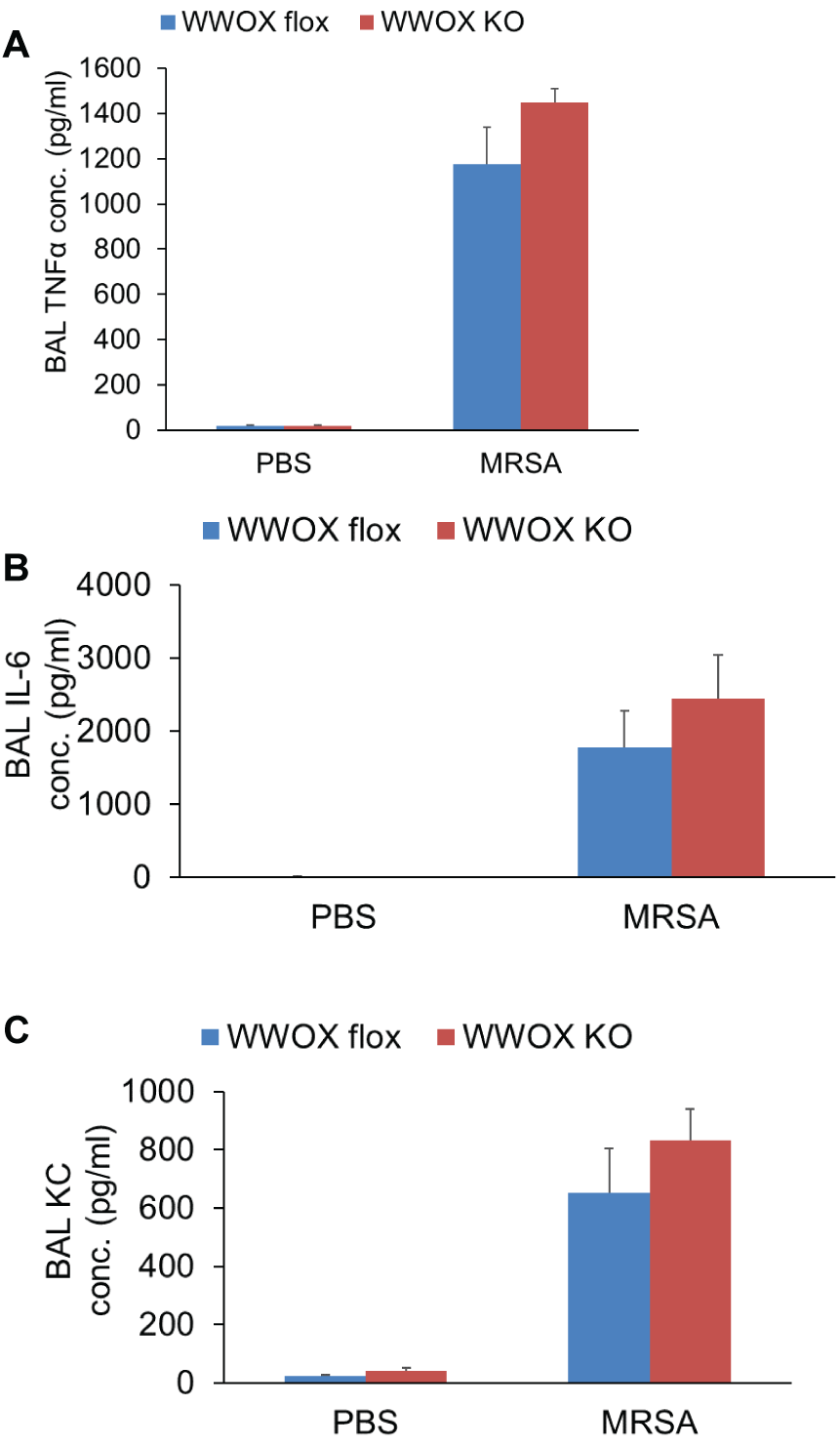
EC-specific WWOX KO mice do not exhibit greater BAL cytokines during MRSA-induced ARDS. Bar graphs depict BAL TNFα, IL-6, and keratinocyte-derived chemokine (KC) concentrations. None of the increases in LPS-treated WWOX KO animals were statistically significantly different than those of LPS-treated controls. Results are shown as means +/- SD in n=3 independent experiments. *p<0.05, compared with LPS-treated control by Student’s t-test.

### ECs isolated from EC WWOX KO mice are more susceptible to MRSA-induced monolayer barrier disruption

ECs from WWOX KO and wild type controls were isolated by flow cytometric sorting for CD31+CD45- cells and grown in culture. They were then examined by ECIS measurement of TER and transwell FITC-dextran passage for their monolayer barrier properties during exposure to MRSA. WWOX KO ECs exhibited a greater decrease in TER during MRSA exposure compared to ECs from wild type animals (**Figure 7A**). The correspondence of this observation to barrier disruption was confirmed by measurement of FITC-dextran passage across monolayers of ECs from WWOX KO mice versus controls in a transwell assay (**Figure 7B**).

**Figure 7.**
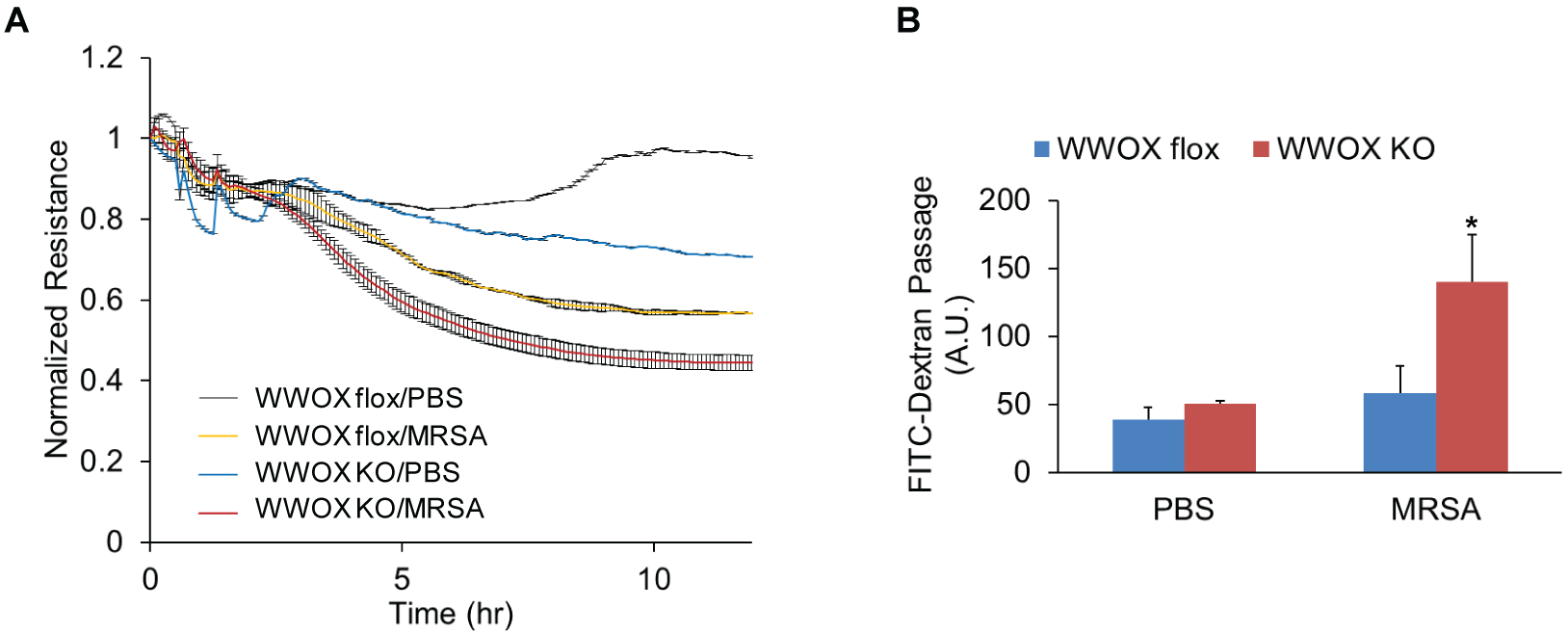
ECs isolated from EC WWOX KO mice are more susceptible to MRSA-induced monolayer barrier disruption. ECs from WWOX KO and wild type controls were isolated by flow cytometric sorting for CD31+CD45- cells and grown in culture. They were then examined by ECIS measurement of transendothelial resistance (TER) during exposure to heat-killed MRSA. **A.** The scatter plot depicts changes in TER over the course of 10 hours following introduction of MRSA to the cell culture media. **B.** In separate experiments, ECs from WWOX KO and wild type controls were grown in a monolayer on transwell inserts containing 3 micron pores. 10 µg/ml FITC-labeled dextran (3kDa) was added over the cells along with MRSA. After 3 hours media from below the transwell was analyzed by fluorometry for the presence of FITC. Results are shown as means +/- SD in n=3 independent experiments. *p<0.05, compared with MRSA-treated control by Student’s t-test.

Cell media supernatants from MRSA-stimulated ECs were examined for concentrations of secreted IL-6, KC, and MCP-1. These were all increased in the supernatants of WWOX KO cells compared to wild types (**Figure 8**).

**Figure 8.**
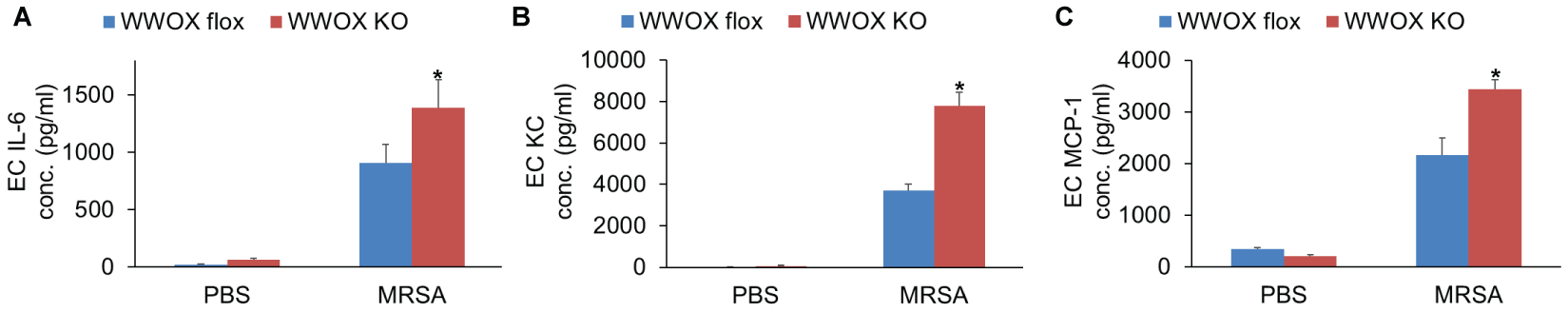
ECs isolated from EC WWOX KO mice produce more cytokines during MRSA exposure. Cell media supernatants from MRSA-stimulated ECs were examined for concentrations of secreted IL-6, KC, and MCP-1. Bar graphs depict these results here. Results are shown as means +/- SD in n=3 independent experiments. *p<0.05, compared with MRSA-treated control by Student’s t-test.

## Discussion

In the current study it has been discovered that loss of lung WWOX expression has additional effects on lung disease susceptibility which were difficult to detect in prior global knockdown experiments that were associated with massive lung neutrophil influx[20]. The current set of experiments focuses on lung vascular endothelium, and suggests that CS/e-cigarette related loss of WWOX expression in these lung cells may underlie the observed increase in susceptibility for severe ARDS seen epidemiologically in chronic smokers[2-4, 6].

We have previously shown that acute whole lung knockdown of WWOX expression in mice causes neutrophilic inflammation[20]. This phenomenon was found to be linked to disinhibition of basal c-Jun activity which upregulated IL-8 (including mouse analogues) expression in alveolar epithelial cells. While exposure to CS or nicotine-containing e-cigarette vapor have both been associated with WWOX downregulation, this is unlikely to occur at such an acutely global scale in humans. It is more likely that loss of human lung WWOX expression occurs heterogeneously over a chronic time course, and therefore any associated neutrophilic inflammation is more likely to be a part of chronic lung disease such as COPD[32].

The mechanistic link between endothelial WWOX downregulation and enhanced LPS-or MRSA-induced vascular leak remains unknown. In the current study, inhibition of JNK did not appear to affect barrier susceptibility during WWOX knockdown (**Figure 1**), suggesting that c-Jun related pathways are not a significant component of the underlying mechanism. The cell supernatant levels of inflammatory cytokines were elevated in MRSA-stimulated WWOX deficient ECs compared to wild types. Some of these may contribute to enhanced EC barrier disruption seen in WWOX-deficient ECs[33] via autocrine effects. However, interestingly, there were no significant differences in the BAL levels of permeability-promoting cytokines between wild type and EC WWOX KO mice treated with MRSA. This observation suggests that additional inflammation-independent mechanisms exist by which EC WWOX deficiency may lead to increased vascular leak.

The association between cigarette smoke exposure and genotoxic stress at the chromosomal fragile site in which WWOX resides in the genome highlights the potential for chromosomal fragile sites to be examined in the context of lung disease. Nonrandom chromosomal aberrations are now known to play a significant role in the development of various human malignancies[34]. Specific chromosomal locations in the human genome that are particularly susceptible to such structural alteration are called fragile sites[34]. These have been further classified broadly into two categories. Those which are thought to be altered relatively rarely, and which have been observed in association with heritable disease, are termed rare chromosomal fragile sites, while those which are altered more frequently in conjunction with toxic environmental exposures are known as common chromosomal fragile sites[34]. The mechanism of chromosomal fragility is thought to be due to structural factors that increase the tendency for gaps or constrictions to form at particular locations which may break during partial replication stress[34]. Rare fragile sites occur only in select individuals, and confer susceptibility to a particular heritable disease[34]. The classic example of a rare fragile site is Fragile X syndrome, a heritable form of intellectual disability[34]. Common chromosomal fragile sites, on the other hand, occur in nearly all individuals and are thought to accrue mutations somatically during genotoxic stress related to environmental exposures[34]. Virtually all common fragile genes have been studied in the context of cancer, where a particular mutation at one of these sites would be perpetuated by additional oncogenic factors. The rate of accumulation in normal tissue is largely unknown. However, interestingly, in at least one study of WWOX expression in cancer, reduced WWOX expression was not only observed in the majority of cancer cells examined, but also in about one-third of adjacent normal tissue[35]. (long but interesting discussion but link to the results needs to be highlighted)

Cigarette smoke has long been known to be one such genotoxic stress that increases the frequency of aberrations observed at common fragile sites[17]. This association has been studied extensively in bone marrow and peripheral blood cells where it is linked to the development of leukemias and lymphomas[17]. However, the accumulation of aberrations at common fragile sites has not yet been studied in the lung where environmental genotoxic stress may theoretically result in the greatest amount of accumulated aberrations over the life of an individual smoker.

While the frequency of fragile site events or the rate of accumulation of chromosomal aberrations in the lungs of smokers is therefore largely unknown, it is interesting to note that WWOX, which resides at the second most active chromosomal fragile site in humans[36], is linked to signaling events relevant for a number of smoking-related lung diseases. WWOX is already known to interact with several molecules linked to the pathogenesis of lung disease including those in which pulmonary vascular leak is a pathophysiologic component. At least eight of the known binding partners of WWOX [36, 37] may connect it to non-cancer-related lung disease. These include NFκB activating protein [38], ErbB4 [39], c-Jun [40-43], ezrin [44], Dvl2 [45], p53 [46, 47], and the R-SMADs [48, 49]. Many of these molecules are already connected to the pathogenesis of lung diseases such as acute respiratory distress syndrome (ARDS), asthma, chronic obstructive pulmonary disease (COPD), pulmonary hypertension (PH), and pulmonary fibrosis (PF) [50-60]. Furthermore, WWOX polymorphisms have been associated with human COPD susceptibility and pulmonary function traits such as forced vital capacity[61-63].

The most critical inference that may be drawn from this study is with regards to the possible long-term effects of e-cigarette use. The mouse model utilizes a dosage and duration that is comparable to human exposures of up to 20 years. Lung WWOX expression was observed to decrease only in mice exposed to vapor containing nicotine. Therefore, this study potentially predicts that the consequences for chronic users of nicotine-containing e-cigarettes would include an increased risk of ARDS during lung infection as well as increased risk for the development of chronic lung diseases which are associated with neutrophilic inflammation. The development of a dual-hit e-cigarette exposure and MRSA-induced ARDS animal model is needed to confirm this association, and to define the relative importance of WWOX downregulation versus other e-cigarette-induced gene expression changes in mediating these potential risks.

## Acknowledgments

This work was supported by the services of the Cardiovascular Research Core at the University of Illinois in Chicago. Funding sources include NIH HL126176 and HL103836 to LBW; R01-HL-127342, R01-HL-133951 2R01 HL111656-06 to RFM; 1K08 HL140222-01A1 to SS.

